# Amber Codon Mutational Scanning and Bioorthogonal PEGylation for Epitope Mapping of Antibody Binding Sites on Human Arginase-1

**DOI:** 10.1101/2024.08.20.608740

**Authors:** Jaime Fernández de Santaella, Nikolaj G. Koch, Lorenz Widmer, Michael A. Nash

## Abstract

Epitope mapping is crucial for understanding immunological responses to protein therapeutics. Here, we combined genetic code expansion and bacterial surface display to incorporate S-allylcysteine (SAC) into human arginase-1 (hArg1) via Methanococcoides *burtonii* pyrrolysyl-tRNA synthetase. Using an amber codon deep mutational scanning and sequencing workflow, we mapped SAC incorporation efficiency across the hArg1 sequence, providing insights into structural and sequence dependencies of non-canonical amino acid incorporation. We used mutually bioorthogonal allyl/tetrazine and azide/DBCO chemistries to achieve site-specific PEGylation and fluorescent labeling of hArg1, revealing insights into SAC side chain reactivity and solvent accessibility of residues in hArg1. This system was further applied to determine the binding epitope of a monoclonal antibody on the surface of hArg1, providing high-resolution data on the impact of PEGylation residue position on antibody binding. Our method produces high dimensional data of non-canonical amino acid incorporation efficiency, site-specific functionalization enabled by mutually bioorthogonal chemistries, and epitope mapping of therapeutic proteins.

## Introduction

Epitope mapping refers to a broad suite of experimental techniques used to identify the specific binding sites or epitopes on the surfaces of antigens that are recognized by antibodies. These techniques are essential for choosing therapeutic antibodies that bind the correct portion of a target and achieve a specific molecular or cellular response. Epitope mapping can further be used to understand the immunological response to protein therapeutics in the context of anti-drug antibodies^1–5^, and help identify immunogenic regions of therapeutic proteins. Once identified, these immunogenic epitopes can be engineered away through sequence alterations that reduce immunogenicity while maintaining activity.

Currently available methods^6^ for epitope mapping include high-resolution structural techniques such as X-ray crystallography^7^ and nuclear magnetic resonance (NMR) spectroscopy,^8^ however, many antibody-antigen complexes do not co-crystallize and NMR may struggle to resolve complexes with high molecular weight (>40 kDa). Cryogenic electron microscopy (cryo-EM)^9^ is an emerging technique, but is limited in terms of resolution, particularly for small epitopes. Mass spectrometry-based methods include hydrogen-deuterium exchange,^10^ cross-linking mass spectrometry,^11,12^ and fast photochemical oxidation,^13,14^ but each have technological or practical limitations.^15^ Peptide microarray methods as well as cell/phage surface display of linear peptides have also been reported, but these approaches do not necessarily represent native antigen structures and may struggle with conformational epitopes.^16–19^

High-throughput epitope mapping by deep mutational scanning (DMS)^20–24^ is a modern method that builds on single variant scanning methodologies such as alanine or cysteine scanning.^25–27^ In these workflows, nearly comprehensive site saturation libraries of the target antigen are generated in a pooled cellular display format and screened by fluorescent activated cell sorting and next-generation DNA sequencing (NGS). This allows experimenters to detect variants that are highly expressed but do not bind the antibody, or bind with decreased affinity.^20,24,28^ By mapping the positions of these mutations onto the antigen structure, the binding epitope can be determined with high resolution.

In this report, we developed an amber codon scanning workflow that leverages advances in genetic code expansion and non-canonical amino acid incorporation. Our approach involves systematically introducing amber (TAG) stop codons within an antigen sequence displayed on *E. coli*, and leveraging genetic code expansion (GCE) with the pyrrolysine system to incorporate the non-natural amino acid S-allylcysteine (SAC).^29^ The bioorthogonal allyl group of SAC^30,31^ is then used for site-specific PEGylation, serving to disrupt access of the antibody to the surface of the antigen at that site. Using PEG as a steric inhibitor of binding offers a method to identify binding epitopes, and to understand how site-specific PEGylation can mitigate recognition by anti-drug antibodies. Incorporation of non-natural amino acids into therapeutic proteins can provide specific bioconjugation points for attachment of drugs of polymer chains^32^, and such techniques have been implemented in bacterial^33,34^, fungal (i.e. yeast)^35^ and mammalian systems. Site-specific PEGylation is commonly used to enhance serum half life and decrease immunogenicity of therapeutic proteins. By generating high dimensional data on the influence of PEGylation on immune recognition, our approach merges epitope mapping with information on optimal PEGylation strategies^36^ for therapeutic formulations.

As a model antigen, we worked with human arginase-1 (hArg1), a pivotal enzyme in the urea cycle and various other metabolic pathways. hArg1 is a homotrimeric manganese-metalloenzyme that detoxifies ammonia in the body by catalyzing the hydrolysis of L-arginine to L-ornithine and urea. This enzyme is a relevant biotherapeutic in the context of autosomal recessive hyperargininemia, which leads to the accumulation of arginine and ammonium ions.^37,38^ The current standard of care includes IV administration of pegzilarginase, a recombinant PEGylated version of hArg1.^39^ Pegzilarginase has further been proposed as a therapeutic agent for treating various types of cancer that exhibit deficiencies in arginine biosynthesis.^40–43^ This approach exploits the metabolic vulnerability of malignant cells that are reliant on extracellular L-arginine. Nonetheless, the effectiveness of these therapies is known to be at least partially limited by the immunogenicity of hArg1. Anti-hArg1 adaptive immune cells have been found in both healthy and cancerous patients.^44–46^ Other factors such as a prolonged exposure to high doses of hArg1, and differences in recombinant protein production/purification methods^47^ may further influence immunogenicity. Our study therefore focused on developing an epitope mapping protocol based on amber codon scanning to identify PEGylation sites within hArg1 that can prevent recognition by anti-drug antibodies.

## Results

### System overview

A summary of our amber codon scanning workflow is shown in **Figure 1**. We employed a two-plasmid system to achieve site-specific incorporation of ncAAs into proteins displayed on *E. coli*. The first plasmid introduced an orthogonal translation system (OTS) that included a constitutively expressed pyrrolysyl-tRNA synthetase (PylRS) and its cognate tRNA. This synthetase loads SAC onto the amber-recognizing tRNA (tRNA^AUC^). When an amber stop codon is encountered during translation, the loaded tRNA incorporates SAC into the peptide chain on the ribosome. The second plasmid contained the displayable hArg1 variant library that was enriched with variants carrying single amber stop codons. Cell-surface display of the hArg1 variant was achieved by its fusion to the N-terminus of the autotransporter AIDA-1.^48^ We previously demonstrated how this fusion construct can successfully display active hArg1 trimers and serve as a tool for screening the activity of hArg1 variants.^49^

**Figure 1:**
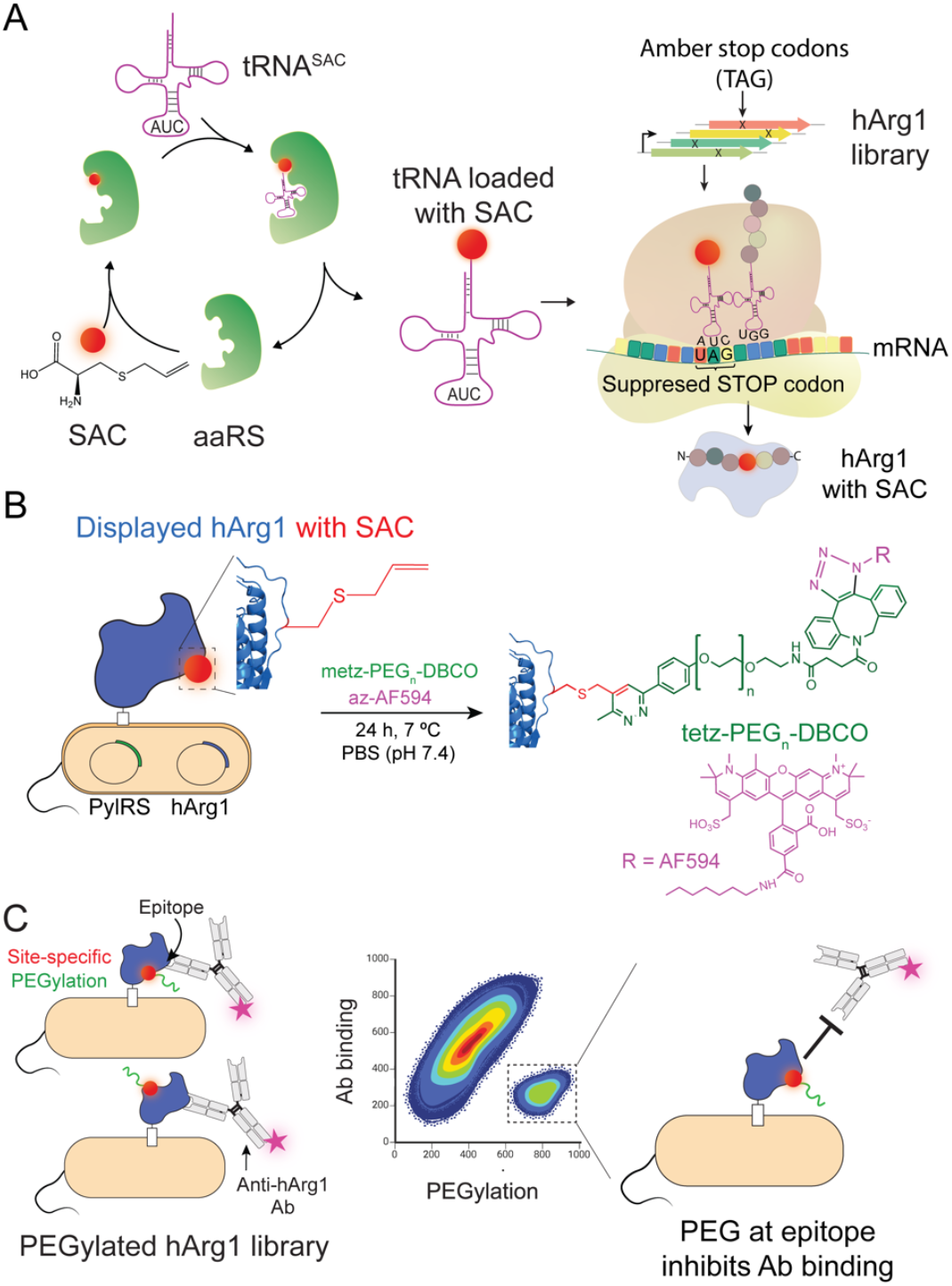
Workflow overview. (A) Orthogonal translation system (OTS) composed of a pyrrolysyl-tRNA synthetase (PylRS - green) tRNA (pink) pair. The PylRS specifically recognizes its cognate tRNA (tRNASAC) and loads the ncAA S-allylcysteine (SAC). The tRNA recognizes amber stop codons and allows translation when the stop codon is reached by incorporating SAC into the nascent polypeptide chain (right). A library of displayed hArg1 variants is first FACS-enriched for amber stop codons and used for site-specific ncAA incorporation into displayed hArg1. (B) Site-specific incorporation of SAC into displayed hArg1 provides a reactive site for conjugation by methyltetrazine-PEG_23_-DBCO and azide-AlexaFluor488 to produce libraries of site-specifically PEGylated fluorescent hArg1. (C) PEGylated displayed hArg1 libraries are panned against anti-hArg1 antibodies. Variants with PEGylated SAC incorporated in the binding epitope will inhibit antibody binding despite high expression levels. Sorting and sequencing of these variants serves to identify the epitope.

We generated the hArg1 variant library in two steps. First, we used the one-pot tiling mutagenesis method with NNK codons to create a ‘mother library’ as is typically done in DMS workflows.^50,51^. The only stop codons encoded by our NNK tiled primers were amber stop codons, which were introduced at nearly each and every position within the hArg1 coding sequence. Second, to enrich variants containing these amber stop codons, we measured hArg1 expression/display levels by antibody staining and flow cytometry, and sorted the variants that did not display hArg1. This cellular pool constituted our library enriched in amber stop codons (i.e. amber scanning library), which was used for subsequent experiments. Nanopore sequencing of the library revealed that ∼85% of all possible single amber substitutions within the hArg1 gene were present. In the presence of SAC, amber stop codons within the hArg1 gene were suppressed and translated with SAC, enabling bioorthogonal site-specific derivatization of surface displayed hArg1 in subsequent steps (**Figure 1A**).

The allyl group of SAC enabled an Inverse Electron-Demand Diels-Alder (IEDDA) reaction,^30,31^ resulting in covalent conjugation of methyltetrazine-(PEG)^n^-DBCO (**Figure 1B**). This click chemistry reaction allowed for efficient bioorthogonal conjugation under physiological conditions, resulting in site-specific PEGylation. Simultaneously, a second mutually orthogonal click reaction was used, where the free DBCO functional group of the bifunctional PEG was conjugated to an azide-fluorophore (**Figure 1B**). As a result, hArg1 molecules that were successfully expressed and PEGylated were detectable by fluorescence.

Once PEGylated, an immunofluorescence FACS assay was used to determine if an anti-hArg1 antibody could bind hArg1 in the presence or absence of conjugated PEG. A common issue in GCE is the occurrence of truncation products caused by unsuccessful stop codon suppression. The AIDA-1 system avoided this issue by fusing the hArg1 variants to the N-terminus of the AIDA-1 surface display anchor. As a result, prematurely truncated hArg1 variants lacked the display anchor. Thus, detection of the PEG-fluorophore indicated the presence of full length hArg1. The intensity of the anti-hArg1 monoclonal antibody binding signal was then correlated with the amount of displayed hArg1. Variants that were highly expressed and PEGylated but showed no antibody binding were then sorted using fluorescence-activated cell sorting (FACS), and the implicated epitope residues subsequently identified through NGS and structural modeling.

#### SAC incorporation efficiency and comparison of PylRS systems

We tested several known PylRS systems, in an AIDA compatible pUltra vector,^52^ to identify one that would incorporate SAC efficiently into hArg1 fused to the N-terminus of an AIDA display anchor^53^. We tested PylRSs from *Methanosarcina mazei*/*barkeri* (MmPylRS/MbPylRS), *Methanococcoides burtonii* (MburPylRS)^54^, and a modified MbPylRS fused with the solubility tag SmbP.55 SAC incorporation efficiency was measured by comparing the level of display of our library using an immunolabeling assay to fluorescently label the N-terminal 6xHisTag of hArg1 (Figure 2A and S1). Since hArg1 lies on the N-terminus of the display anchor, any stop codon not subject to amber suppression will result in no signal. This means that the hArg1-AIDA fusion system that we tested here is built in such a way that hArg1 along with the N-terminal 6xHisTag is only displayed when the C-terminal autotransporter sequence is successfully translated. Any premature termination within the hArg1 sequence will therefore lead to no surface display, and no cell labeling through the 6xHisTag. To phrase it differently, only full-length sequences with successfully suppressed amber stop codons are capable of being displayed.

**Figure 2:**
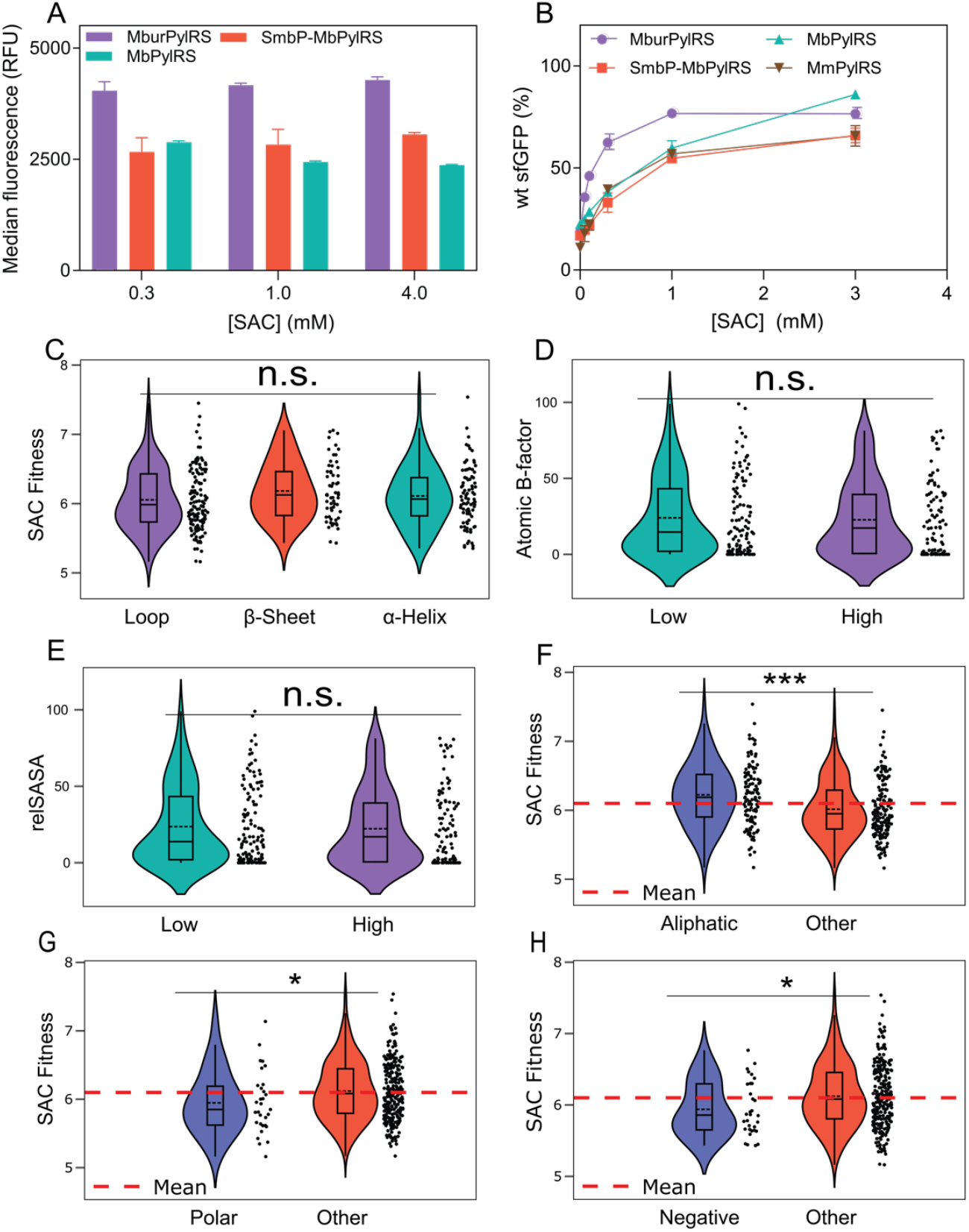
PylRS comparison and SAC incorporation analysis. (A) Immunofluorescence assay characterizing hArg1 display level for an amber-rich library with different PylRS systems. Median fluorescence of the displaying cells measured by flow cytometry reports on SAC incorporation efficiency. For cells cultured in the absence of SAC, few to no events were detected in the cytometer gate (see Figure S1) (median ± SD, n = 3). (B) Comparison of different PylRS/SAC incorporation efficiencies on soluble single amber stop codon variants of sfGFP. Assay performed on bacterial strain B-95.ΔAΔfabR^57^ (see Figure S2). (C-F) Investigation of the dependence of MburPylRS SAC incorporation efficiency on the secondary structure properties of the substituted amber codon site. Effect of secondary structure classification (C), atomic B-factor (D), per residue solvent-accessible surface area (E) and residue class, including aliphatic (F), polar (G) and negative (H) residues. SAC Fitness was categorized as ‘High’ or ‘Low’ if their values were further than 1 standard deviation from the mean. A normality test was performed between the populations, and when the population distribution was normal an ANOVA test was performed. For non-normal distributions we performed a KS-test. (n.s.): not significant i.e. p-value > 0.05; (*) p-value < 0.05; (***) p-value < 0.001.

Using this N-terminal 6xHisTag as a readout, we found that MburPylRS outperformed the other PylRSs across all SAC concentrations. We used MburPylRS with 4 mM SAC for the remainder of the work. Surprisingly, using MmPylRS failed to display hArg1 even though activity of this PylRS was clearly observed using an amber-containing soluble superfolder GFP (sfGFP) assay (data not shown). Otherwise, the display efficiencies of hArg1 tested with the different OTSs correlated with sfGFP expression assays (Figure 2B and S2).

We next analyzed the amber codon scanning library, which introduced single in-frame amber-stop codons across the coding region of hArg1. This method allows for comparing incorporation efficiency of different OTSs or different ncAAs by FACS sorting according to display levels followed by NGS and variant scoring. This procedure generates expression fitness scores that allow comparison of ncAA incorporation efficiency as a function of substituted residue position within the sequence/structure of hArg1. Additionally, the amber scanning methodology can be used to sort, sequence and score variants based on their ability to react with the given ncAA, for example by direct labeling or PEGylation (see below).

We first applied this NGS-based variant scoring methodology to characterize site-specific SAC incorporation, which we call SAC fitness score. We quantified expression levels for variants contained in the amber codon enriched hArg1 library by fluorescently immunolabeling the N-terminal 6xHisTag fused to hArg1. Since the library was first enriched for non-displaying mutants, hArg1 library display levels in the absence of SAC were minimal (Figure S1). When SAC was provided to such cells, the incorporation of SAC into the sequence led to amber suppression and generated substantially higher display levels. Cells were sorted into four bins based on expression levels and analyzed using NGS. Mutants were sorted into a non-expressing population (Bin 1), and into three further bins (Bins 2-4) according to increasing fluorescence intensity, with the aim of achieving a comparable proportion of expressing cells within each of these latter bins. The SAC fitness score was calculated from the NGS reads for each variant based on its relative abundance across these bins. This value then serves as a relative measure of site-specific SAC incorporation efficiency. To calculate SAC fitness score, the number of NGS reads per variant was first converted into a number of cells (Supporting Information, Eq. 2). This cell number together with the median fluorescence value per bin was then used in a weighted average calculation to quantify the mean fluorescence per variant (Supporting Information, Eq. 3). The log^10^ value of this mean served as the SAC fitness score (Supporting Information, Eq. 4 and Eq. 5 -see Supporting Information for details on fitness score calculations).

We calculated linear correlations of fitness scores determined by NGS for biological replicates, and found a correlation coefficient of 0.91, demonstrating high reproducibility of the method (Figure S3A). A comparison of fitness scores between amber-containing single mutants and other mutations revealed significantly higher scores for the former (Figure S3B). This indicates that variants containing amber stop codons were expressed at a higher level than those lacking stop codons. This finding was attributed to having pre-sorted and pre-enriched the mother library for non-expressing variants in the absence of SAC, resulting in a library enriched with sense mutations that hinder the display hArg1 as well as stop codons that generate no expression signals. Stop codons within the hArg1 gene result in no expression because the AIDA display anchor is at the C-terminal end of the construct. The NGS data were used to generate expression fitness scores for each amber single-mutant variant in the library, creating a site-specific map of SAC incorporation and display efficiency (Figure S3C). Our library included single amber-containing variants in 85% of the hArg1 sequence positions (274/322 residues). Among these, 124 variants (45%) demonstrated significantly higher SAC incorporation scores compared to the non-amber-containing mutants, which served as a negative control (p-value < 0.05, KS-test).

Further analysis revealed that the incorporation of SAC was not dependent on secondary structure (Figure 2C), atomic B factor (Figure 2D), or relative solvent accessible surface availability (relSASA) of the side chain (Figure 2E). Here, SAC Fitness was categorized as ‘High’ or ‘Low’ if their values were further than 1 standard deviation from the mean and these groups were then compared. However, for certain biophysical properties of the substituted residue, we did observe a significant influence on the SAC-incorporation (i.e. expression level) scores. For instance, polar and negatively charged residues were significantly less likely to be successfully substituted by SAC than any other residue, especially when compared to aliphatic residues, which had the highest SAC incorporation efficiencies (Figure 2F - 2H). We also found that the nucleotide context of the incorporation site had a significant influence on the SAC incorporation scores. For example, if the nucleotide directly downstream (position n + 1) from the amber-stop codon was a cytosine, the SAC fitness score was more likely to be higher than for guanine (Figure S4). Other codon dependencies were found and described in Figure S4. Efforts to characterize the effect of mRNA context on ncAA incorporation have thus far been limited, however, prior work has shown that for specific OTS and ncAA pairs, examples are transferable from protein to protein.^56^ Thus, producing these databases with general nucleotide preferences can be valuable and potentially generalizable to other targets. Interestingly, we find that nucleotides directly downstream of the amber stop codon have a larger impact than those upstream of it (Figure S4). To our knowledge, this work represents the first report combining genetic code expansion with bacterial protein display. By implementing high-throughput NGS-based expression analysis of the amber scanning library, we obtained high resolution site-specific data on SAC incorporation efficiency. This analysis reveals several biophysical factors of the amber codon insertion position that influenced SAC incorporation efficiency.

#### Bioorthogonal site-specific PEGylation

Next we investigated bioorthogonal click reactions to attach fluorophores and polymers to SAC incorporated into hArg1 on the bacterial cell surface. We performed a fluorescent labeling experiment with pyrimidyl-tetrazine-AF488 (Py-Fluor) and characterized the reactivity by flow cytometry. Successful bioconjugation of the fluorophore occurred only on cells that had been grown with 4 mM of SAC. To optimize this reaction, various times, temperatures, SAC concentrations and Py-Fluor/cell ratios were tested (Figures S5A & S5B). The best condition was found to be a 22-hour reaction at 7 °C with 15 μM of Py-Fluor and cells grown with 4 mM SAC (Figure S5C).

We applied the same bioorthogonal conjugation approach for site-specific PEGylation. We used a bifunctional methyltetrazine-mPEG_n_-DBCO (metz-mPEG_n_-DBCO) to PEGylate hArg1 at the position of incorporated SAC, and further derivatized the PEG with an azide-fluorophore (azide-AlexaFluor 594) to enable detection by flow cytometry (Figure 1B). Due to the mutually bioorthogonal nature of both click chemistry reactions (allyl/tetrazine and azide/DBCO), the two reactions could be performed simultaneously in one pot by simply incubating all components in PBS.^58^ We performed the reaction at low temperature (7 °C) to inhibit bacterial growth during the reaction. An optimization experiment analogous to the Py-Fluor reaction was first performed (Figure S5C). In addition, we tested a range of PEG lengths, ranging from 7 to 23 ethylene oxide monomer units, as well as a tri-branched dendrimeric PEG ((methyltetrazine-PEG_10_)-tri-(azide-PEG_10_-ethoxymethyl)-methane) (Figure S7). We detected successful PEGylation and subsequent fluorophore derivatization under all conditions, with the highest labeling occurring at 7 °C following 40 hours of reaction with an excess of fluorophore to PEG of 2:1 (Figure S5).

Similar to the experiments measuring SAC incorporation efficiency, we used gated FACS sorting and NGS to produce both Py-Fluor and PEGylation scores for each SAC insertion position. We generated fitness scores (i.e., Py-Fluor and PEGylation fitness scores) with the same binned sorting/sequencing and variant scoring methodology as described for the SAC incorporation fitness score (Figure 3). For both these cases, instead of assigning bins according to antibody labeling, they were assigned based on increasing Py-Fluor or PEG labeling intensity. Similar controls for reproducibility and separation of non-amber-containing variants were used to validate these datasets (Figure 3A - 3D). In analyzing the raw data, we found a significant correlation between the fitness scores of the click reaction (PEGylation/tetrazine fluorophore fitness scores) and SAC incorporation efficiency (Figures 3F & 3G). Higher SAC fitness scores led to higher PEGylation/tetrazine fluorophore reaction scores consistently across different biological and experimental conditions. This was further supported by the strong correlation and lowest variance between the PEG and Py-Fluor scores. This indicated that the determining factor for both labeling reactions was the presence of SAC to participate in the IEDDA reaction (Figure 3H).

**Figure 3:**
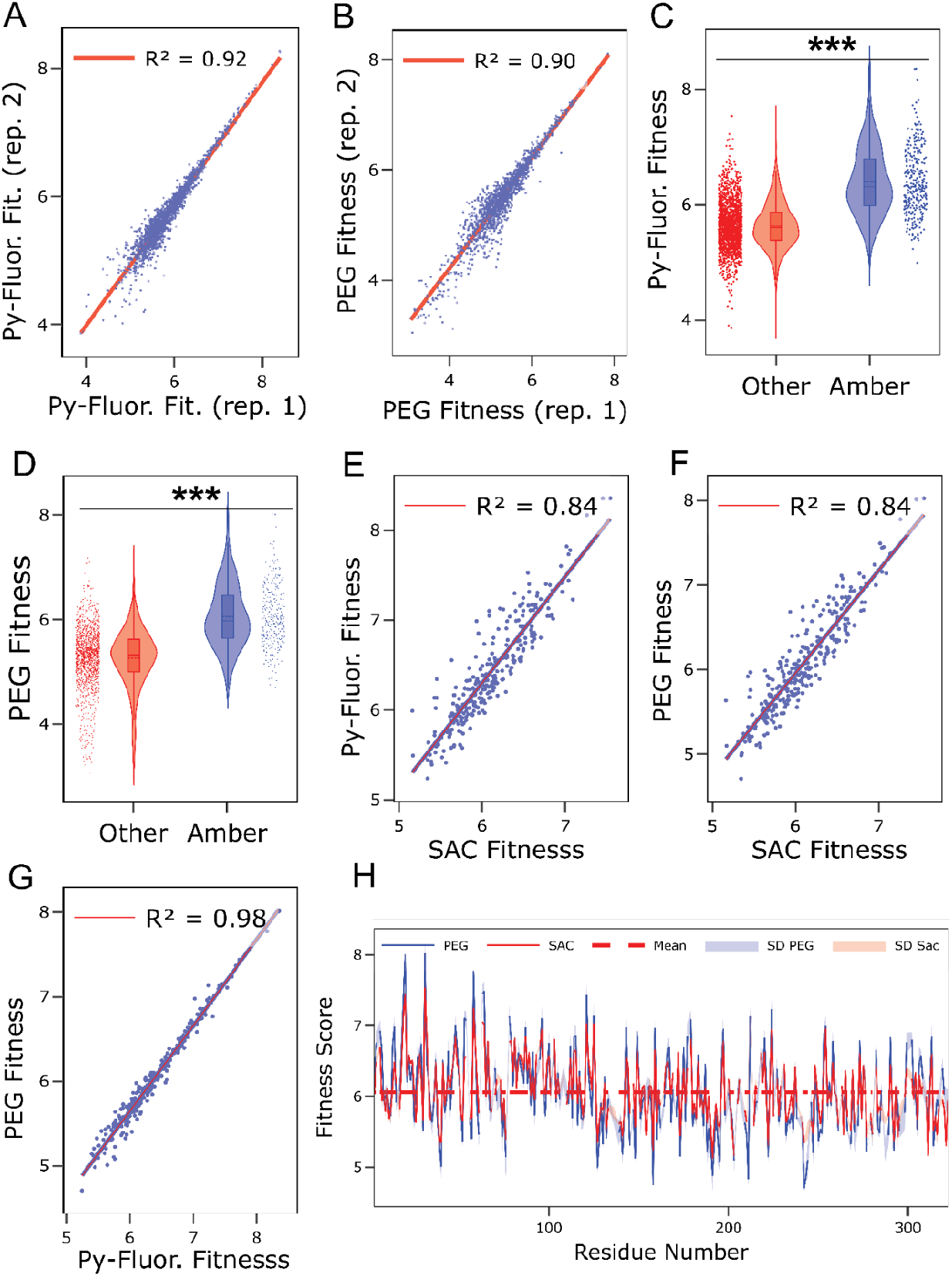
Bioorthogonal reactions performed on mutational amber-scanning library of hArg1. (A) Reproducibility assessment of NGS-derived Py-Fluor fitness scores generated by reacting the amber-scanned hArg1 library with pyrimidyl-tetrazine-AF488. (B) Reproducibility assessment of NGS-derived PEGylation fitness scores generated by reacting the amber-scanned hArg1 library with metz-mPEGn-DBCO/azide-AF594. (C) Distribution of Py-Fluor fitness scores for variants containing amber stop codons compared to variants containing other NNK codon mutations. (D) Distribution of PEGylation fitness scores for variants containing amber stop codons in comparison to variants containing other NNK mutations. Cross-correlation of combinations of three NGS-derived fitness scores across different experiments, (E) SAC incorporation with Py-Fluor fitness score, (F) SAC incorporation with PEG fitness score and (G) Py-Fluor with PEG fitness score. (H) NGS-derived fitness scores for SAC incorporation (red) and PEGylation (blue) as a function of residue number. The standard deviation (SD) is shown as shading. The mean fitness of all amber-containing mutants is shown as a dashed red line. (***, p < 0.001 for both Student t-test and Kolmogorov-Smirnov test).

To investigate the underlying principles governing PEGylation scores, we then normalized the PEG fitness score per residue to the respective SAC incorporation score. This served as a control for hArg1 display level and for the quantity of allyl moieties available for functionalization on the outer *E. coli* membrane. We categorized a PEGylation score as “High PEG_norm_” if it was more than one standard deviation above the mean. Similarly, we categorized as “Low PEG_norm_” those variants that were more than one standard deviation below the mean (Figure S6A). Subsequently, we compared the relSASA of High and Low PEG_norm_ populations, and found that variants classified as high PEG_norm_ exhibited significantly higher relSASA values (Figure S6B), indicating that surface accessibility was associated with a higher degree of PEGylation. Interestingly, some variants with high PEG_norm_ scores exhibited relSASA values of zero, which corresponds to fully buried residues. This suggests that over the 40 hour reaction period, some of the hArg1 residues with relSASA of zero were at some point exposed to solvent and able to react. RelSASA values were calculated from static x-ray crystal structures and thus do not account for dynamics within the structure. We further mapped the positions exhibiting High PEG_norm_ score and relSASA values of zero, and found that they group together in patches (Figure S6C).

#### Epitope Mapping

Finally, we applied this system for site-specific PEGylation and antibody epitope mapping. We bioconjugated hArg1 with the longer PEG (metz-PEG_23_-DBCO), and performed an immunolabeling assay with a commercial anti-hArg1 antibody. Cells that exhibited PEGylation but no antibody binding were sorted and sequenced in Bin 1 (Figure 4A). To identify variants involved in antibody binding inhibition, we calculated a normalized cell number per variant and bin (*C*_*vi*_), as described above (See Supporting Information, Eq. 2). Then, the abundance of each *C*_*vi*_ was compared across bins to assign an abundance score (*Ab*_bin 1_) (See Supporting Information, Eq. 6). The same was done for cells which showed PEGylation and antibody binding in Bin 2. These abundance scores were reproducible across biological replicates (R^2^ = 0.76, Figure 4B). Finally, we compared the abundance score of each bin and selected those which were highly abundant in Bin 1 (*Ab*_*bin* 1_ > *Ab*_*bin* 1_ + (1.5 × *SD*_*bin* 1_)), where *Ab*_*bin* 1_ is the mean cell abundance in Bin 1. Moreover, we controlled for highly occurring Bin 2 variants to reduce the likelihood of false positives, due to the wider fluorescence distribution. Thus, very highly abundant variants from Bin 2 (*Ab*_*bin* 2_ > *Ab*_*bin* 2_ + (2 × *SD*_*bin* 2_) were excluded (Figure 4C). The control against Bin 2 reduces the likelihood of false positives that arise from “leaky” high abundant variants that during flow cytometry get assigned to a low antibody binding score due to their wider distribution.

**Figure 4:**
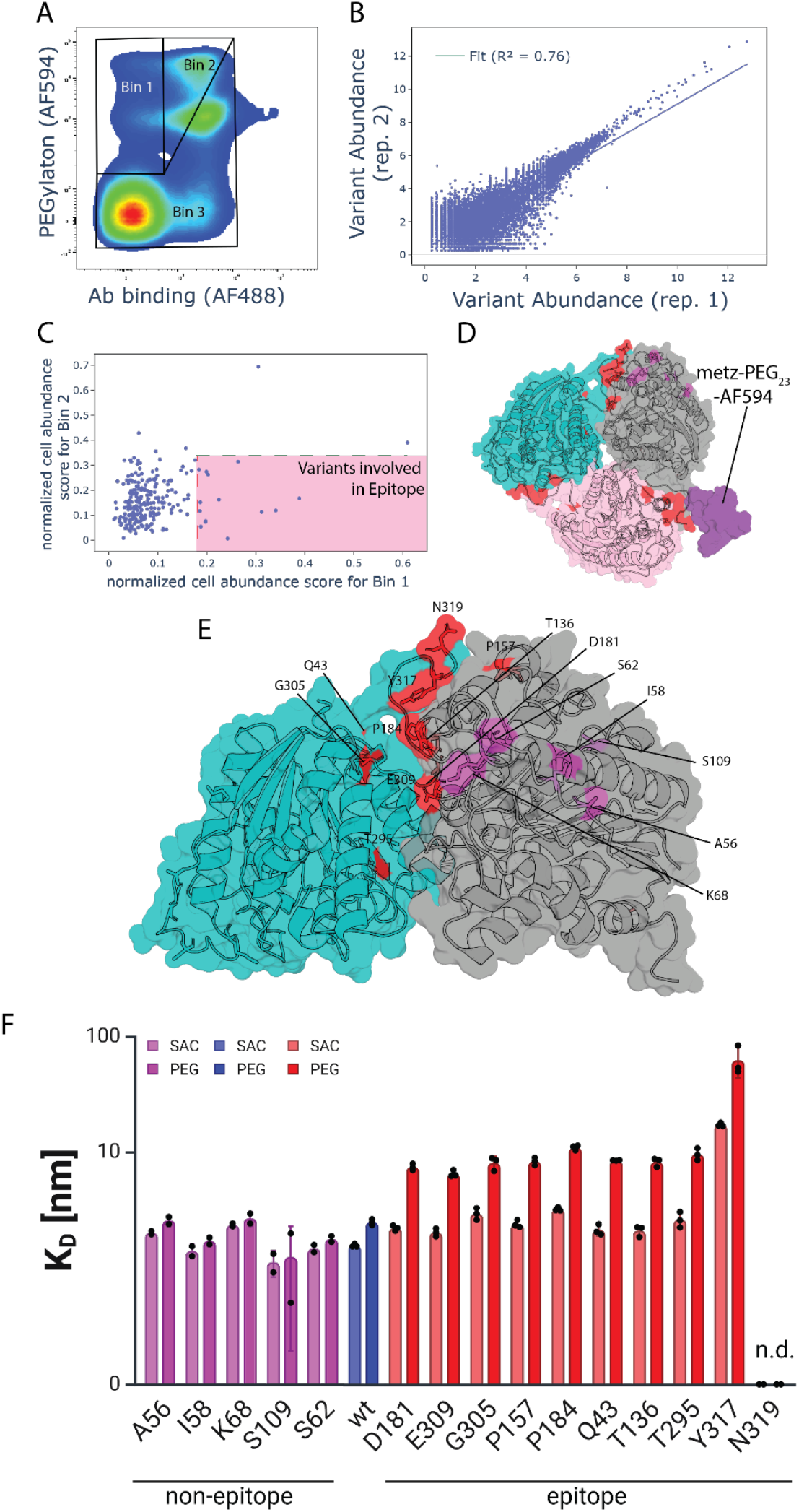
Epitope mapping of an antibody binding site on hArg1. (A) FACS-based immunofluorescence assay and binding assessment using amber-scanning PEGylated hArg1 library displayed on *E. coli* is used to identify and sort variants that inhibit antibody binding. (B) Reproducibility of the number of normalized cells for each variant between two biological replicates. (C) Relative variant abundance (*Ab*_bin 1,_ *Ab*_bin 2_) per bin. Variants highly enriched in Bin 1 and with low abundance in Bin 2 were identified to determine the epitope. (D) Cartoon and surface representation of the hArg1 homotrimer (PDB: 6Q92) with monomers represented in pink, gray and teal. Positions implicated in the antibody binding epitope are colored in red and positions away from the epitope are in magenta. A fold/structure prediction for metz-PEG_23_-azAF594 is shown to scale in purple. (E) Detailed view of non-epitope residues (magenta) and proposed epitope residues (red) located at the interface of two hArg1 monomers. F) Dissociation constants (K_D_) tested in monogenic assays for epitope (red) and non-epitope (magenta) residues, where either only SAC was incorporated (lighter color) or SAC was also PEGylated (bolder color). Biological replicates for epitope residues n=3 and for non-epitope residues n=2; n.d. = not detected. A table with the raw K_D_ data is in the supplements. Significance was tested with a one-way-ANOVA Brown–Forsythe test with a consecutive Dunnett’s T3 multiple comparisons test, for the wt in comparison to the SAC and PEG constructs (see Table S1).

The selected population, highlighted in Figure 4C, contains hArg1 variants bearing a SAC residue at positions that inhibit antibody binding when PEGylated. This population includes variants incorporating SAC within the antibody epitope and other proximal sites where PEGylation sterically inhibits binding. The residues which were PEGylated in these variants are highlighted in red on the crystal structure of the trimeric form of hArg1 (Figure 4D & 4E). The implicated positions were localized proximal to the flexible loop at the C-terminus, a site recognized by other commercial antibodies (e.g. as stated in the specifications of other commercial anti-hArg1 antibodies, Invitrogen MA5-36060).

Interestingly, the loop of hArg1 identified as part of the antibody binding epitope is partially involved in trimerization, and each C-terminus is heavily in contact with another monomer. Our results therefore show that residues from both monomers are involved in the epitope. Considering that we employed a large PEG molecule (depicted to scale in Figure 4D), some of these residues could be epitope adjacent (e.g., Q43, T295, G305) but not directly participate in antibody binding, and/or generally destabilize folding in this region.

The epitope hits from the pooled sequencing experiments were next produced as monogenic variants displayed on *E. coli* and tested following SAC incorporation and PEGylation using a FACS-based antibody binding assay. The resulting antibody binding data (Figure 4F) confirmed most of the hits (statistical analysis presented in Table S1). PEGylation at residue Y317 increased the detectable K_D_ the most, while for N319 no antibody binding was observed. These monogenic epitope validation experiments (Figure 4F) also imply that the incorporation of a site-specific SAC and/or PEG offer a method to tune the sensitivity of our method. By precisely functionalizing hArg1 with SAC or PEG at specific positions, we can amplify the perturbation at the antibody binding interface with larger binding interference occuring with PEG. Furthermore, modulating the size of the PEG molecule could allow for a tunable approach in the future, facilitating a balance between sensitivity and precision in epitope mapping.

## Conclusion

In summary, our study introduces a novel method combining deep mutational scanning, genetic code expansion and bacterial display to provide insights into the reactivity of bioorthogonal functional groups within hArg1 and serve as a platform for epitope mapping of anti-drug antibodies. The combination of site-specific S-allylcysteine incorporation, bacterial cell surface display of hArg1 and the use of mutually bioorthogonal chemistries generates high-throughput data at single amino acid resolution. Following incorporation of S-allylcysteine, we harnessed its bioorthogonal properties to enable site-specific derivatization with small molecule fluorophores and PEG polymer chains. Taken together, we showed which positions in hArg1 preferentially accept S-allylcysteine substitutions, produced detailed insights into the structural and functional dynamics of hArg1 and its bioconjugation, and established a robust platform for mapping the binding sites of antibodies to hArg1.

## Acknowledgments

This work was supported by the University of Basel, ETH Zurich and a Consolidator Grant from the Swiss State Secretariat for Education, Research and Innovation (SERI) to MN.

## Data Availability Statement

The raw data associated with this work are available on the Zenodo repository (DOI: *[will be inserted prior to publication]*)

## Supporting Information

Additional experimental details (Materials and Methods) and expanded datasets including raw data of SAC incorporation efficicnecy (Figures S1 & S2), SAC amber-scanning validation data (Figure S3), data on the incorporation efficiency of SAC by nucleotide context (Figure S4), Fluorophore and PEG reaction conditions optimization (Figure S5), Dynamics of hArg1 studied by PEG amber-scanning (Figure S6), additional PEG bioconjugation data (Figure S7), and statistical analysis of binding affinity of selected variants (Table S1).

## TOC Graphic

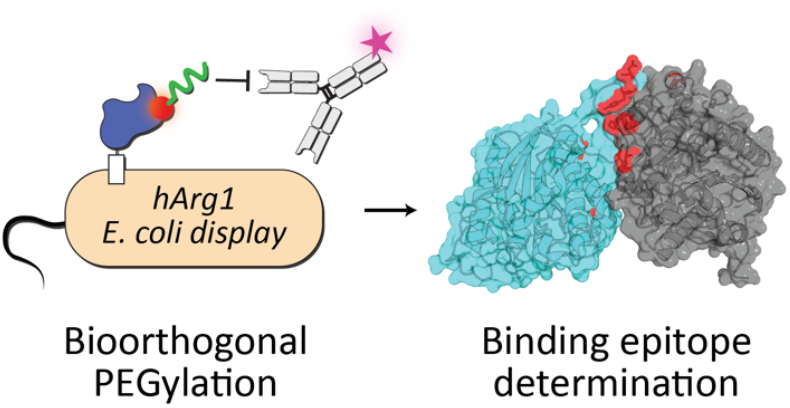

