## Supplementary Information for "Amber Codon Mutational Scanning and Bioorthogonal PEGylation for Epitope Mapping of Antibody Binding Sites on Human Arginase-1"

### Materials and Methods

**hArg1 Cloning and Display.**

The gene coding for hArg1, codon-optimized for expression in *E. coli*, was purchased from Twist Bioscience. The sequence was cloned via Gibson assembly into a pAIDA-1 plasmid, a bacterial surface display vector with a chloramphenicol resistance gene. The hArg1 gene was fused to the N-terminus of the AIDA autotransporter with a glycine-serine linker, and a 6xHisTag was included between the signal peptide and hArg1 for immunolabeling. After sequence confirmation through Sanger sequencing, the plasmid pAIDA1_hArg1 was chemically transformed into the bacterial strain BL21 (DE3), and positive colonies were selected on LB agar with chloramphenicol. Display and activity were confirmed as previously described.[1]

**Orthogonal Translation System Plasmid Construction.**

Since the pAIDA-hArg1 display system had a Chloramphenicol resistance gene and a p15A origin of replication, exactly as the established PylRS OTSs,[2] the different variants were transferred into a system which was compatible, the pUltra vector. The pUltra is an established OTS vector and contains the CloDF13 (CDF) origin of replication and a Spectinomycin resistance marker.[3] We transferred the complete sequence, starting upstream of the PylRS at the lpp promoter and ending at the $tRNA^{Pyl}$ proK terminator, to the pUltra vector. Afterwards we deleted the lac repressor (LacI) gene from the original backbone. All PylRSs contained the C313W:W382T mutations (*M. barkeri* notation) to recognize SAC. All cloning steps were done by golden gate cloning and confirmed by sanger and full plasmid sequencing. The full plasmid sequences can be downloaded from the supporting information.

**DNA Synthesis of DMS Library.**

To produce the hArg1 variant library, the gene was inserted into a pUC19 cloning plasmid. A deep mutational scanning (DMS) library was generated by implementing the plasmid-based one-pot nicking mutagenesis strategy.[4,5] In short, one strand of the hArg1 plasmid was digested using strand-specific DNA nickases (Nt.BbvCI and Nb.BbvCI) and exonuclease activity. The remaining strand then served as a template in a mutagenesis PCR amplification, where a library of tiled 5'-phosphorylated NNK-containing primers was introduced to scan every codon of hArg1 and produced in-frame NNK residue substitutions. A low primer-to-plasmid was used to favor the production of single substitution mutants during the PCR amplification. Subsequently, we digested the template strand using the same nickase-exonuclease strategy and amplified the new variant strands using whole plasmid PCR amplification. The process was repeated again to minimize the percentage of wild-type mutants, resulting in a library size of over 800,000 individual colonies. To reduce complexity, 150,000 cells were counted and sorted by flow cytometry, forming our stock DMS library for ongoing studies of hArg1 variant stability and activity. For bacterial cell-surface display, the hArg1 gene, flanked by a 6x-HisTag, was integrated into the N-terminal terminus of an AIDA-1 autotransporter anchor protein and transformed into *E. coli* BL21 (DE3).[1]

**Generation of Amber Scanning Library by Stop Codon Enrichment.**

The use of NNK codons during the synthesis of the mother library meant that variants containing amber stop codons (TAG) were frequent members of this library. We performed an immunofluorescence staining and FACS sorting experiment to enrich variants that were unable to display hArg1 due to the presence of an amber stop codon within the hArg1 coding sequence. Cells were cultured for 24 hours at 37°C with 0.5 mM IPTG in LB media, washed twice with phosphate-buffered saline with 0.1% bovine serum albumin at pH 7.4 (PBS-BSA), and incubated with a 1:500 dilution of a mouse anti-6xHis antibody (Invitrogen, MA1-21315) in PBS-BSA for 30 minutes at room temperature. After two additional PBS-BSA washes, cells were incubated in ice with a 1:500 dilution of goat anti-mouse IgG conjugated with an Alexa Fluor 594 (Invitrogen, A-11005) in PBS-BSA. Following two final PBS washes, cell sorting was performed using a BD FACSMelody sorter. $10^6$ non-displaying cells were sorted and regrown overnight in LB media at 37°C, enriching the mother library for amber stop codons while retaining non-displaying hArg1 mutants. The DNA from these sorted cells was mini-prepped and co-transformed with the corresponding OTS system into BL21 (DE3) cells. Approximately 5,000 colonies formed the amber-codon enriched scanning library, which was stored at - 80 °C with 25% (v/v) glycerol.

**OTS Efficiency Comparison of SAC Incorporation into Displayed hArg1.**

SAC incorporation in displayed hArg1 was achieved by culturing co-transformed BL21 cells in ZYP-5052 autoinduction media. $OD_{600}$ nm was normalized to 0.05 from overnight pre-cultures grown in LB from the cryostock. ZYP media was supplemented with 4 mM SAC if not otherwise specified. SAC incorporation was assessed by comparing the basal expression level of the library (grown with 0 mM SAC) to those which had soluble SAC present in the media. Expression level was evaluated with an immunofluorescence assay combined with flow cytometry identical to the one described above. In this case, median fluorescence of the positively-gated population was obtained. ±SD (n = 3 biological replicates).

**OTS efficiency comparison of SAC incorporation by sfGFP intact cell fluorescence.**

To quantify the performance of the different PylRS OTSs in the pUltra backbone, an established intact cell fluorescence sfGFP expression assay was performed (Figure S2). *E. coli* B-95.ΔA.ΔfabR (Addgene #197934) cells were used for the reporter assay. Chemically competent cells were first transformed with the pET-28a (SUMO-sfGFP) reporter constructs containing either no stop codon (wild-type) or containing 1x, 3x or 5x amber codons. A second round transformation with chemically competent cells already containing the reporter plasmids introduced the desired orthogonal translation systems and yielded the final cells. LB agar plates for plating contained 1% glucose and corresponding antibiotics. Single colonies of clones were used to inoculate 2 mL of LB medium (in 14 mL tubes) with 1% glucose and appropriate antibiotics, and cultures were grown to saturation overnight. Assays were conducted in 96-well plate format. Cultures were added to each well at 1:100 dilution in ZYP 5052 auto-induction medium to a final volume of 100 µL, supplemented with antibiotics and SAC. Cells were grown in black µ-plates with a clear bottom (Greiner Bio-One, Kremsmünster, Austria, Item No.: 655097) covered with a gas permeable foil (Breathe-Easy®, Diversified Biotech, Dedham, MA, USA) with orbital shaking (Thermo Scientific MaxQ 4450) for 24 h, 300 rpm at 37 °C. For plate reader (Tecan Safire 2, Männedorf, Switzerland) endpoint measurements, the plate foil was removed before fluorescence and $OD_{600}$ measurements. Excitation and emission wavelengths for fluorescence measurements were set to 485 nm and 510 nm, respectively. Fluorescence was normalized by the corresponding $OD_{600}$ values. Biological triplicates were used for measurements of each PylRS construct. Then the relative fluorescence was normalized to the signals of wild-type sfGFP containing no stop codons. ±SD (n = 3)

**Click Chemistry Fluorophore Labeling and PEGylation Reaction.**

For fluorophore labeling and PEGylation, SAC was incorporated as described above: in autoinduction media supplemented with 4 mM SAC. For fluorophore labeling, unless otherwise stated, 200 million cells were washed twice in PBS and incubated with 15 µM of pyrimidyl-tetrazine-AF488 (Jena Bioscience, CLK-103) for 22 hours at 7 ºC with shaking at 600 rpm in 100 µL PBS. For PEGylation, unless otherwise stated, 200 million cells were incubated with 15 µM methyltetrazine-$PEG_4$-DBCO (BroadPharm, BP-25740) and 30 µM of azide-AF594 (Jena Bioscience, CLK-1295-1) for 24 hours at 7 ºC and shaking at 600 rpm in 100 µL PBS. Other PEG molecules were also successfully bioconjugated to hArg1, including methyltetrazine-$PEG_7$-DBCO (BroadPharm, BP-25742), methyltetrazine-$PEG_{23}$-DBCO (BroadPharm, BP-25743), and methyltetrazine-PEG10)-Tri-(Azide-PEG10-ethoxymethyl)- methane (BroadPharm, BP-25705) under the same conditions. After the incubation, cells were washed twice with PBS-BSA with 0.05% Tween20 (PBS-BSA-T20) and then twice with PBS-BSA before performing an immunofluorescence labeling assay equivalent to that described above in the OTS comparison experiment. Labeled cells were kept at 4 ºC until analyzed by flow cytometry in a BD FACSMelody.

**Flow Cytometry and DNA Purification for Amber-Scanning Analysis.**

To produce a fitness score for SAC incorporation or click chemistry, cells were binned according to the relevant fluorescence intensity. The negative gate (with the lowest fluorescence signal) was generated using a negative control, which consisted of cells cultured without SAC. The next three gates were populated with an approximately equivalent number of cells each with an increasing fluorescence signal. At least 700,000 cells were sorted per replicate and regrown at 37 ºC in LB with the appropriate antibiotics. From these cells, DNA was purified using the GeneJET Plasmid Miniprep Kit (Thermo Fisher, K0502), and cleaned using the Clean & Concentrator-5 Kit (ZymoPrep, D4013) by eluting with 20 uL of Milli-Q water. The DNA was then stored at -20 ºC until further processed.

**Next-Generation Sequencing by Oxford Nanopore Technologies (ONT) sequencing.**

DNA concentration was quantified with Qubit fluorometric quantification, and DNA was purified with AmpPure XP beads (Beckman). For targeted amplicon sequencing with reduced amplification bias, the hArg1 gene was extracted by performing a double restriction digest with 20 units of EcoRV-HF and of XbaI in rCutSmart buffer at 37 ºC for 1 hour. Libraries were prepared and indexed following the protocol for ligation sequencing amplicons from the Native Barcoding Kit 24 V14 (ONT, SQK-NBD114.24). In short, DNA was first repaired and prepped for ligation using an end-prep kit (NEB, E7546). Then, barcodes and sequencing adapters were sequentially ligated to it before loading onto a MinION Flow Cell (R10.4.1). Sequencing was run for 72 hours resulting in 10 million reads. Basecalling was performed on Dorado (version 0.5.2+7969fab) using the dna_r10.4.1_e8.2_400bps_sup@v4.3.0 basecalling model.

**Variant Calling.**

The resulting reads were processed and filtered for quality, retaining only those with a mean quality score greater than 15. To identify the mutated codons, we used a custom script. In short the reads were aligned to the wild-type hArg1 gene using minimap2 (version 2.26-r1175), and any reads that did not fully map to the reference gene were discarded. Codons that differed by at least one base from the wild-type sequence were identified using samtools pileup (version 1.19). Codons containing indels were considered sequencing errors and were excluded from further analysis. The identified codon mutations were then filtered based on their prevalence in the entire sequenced library and the mean of the lowest basecaller certainty of any base within each codon (as described in Equation 1).

$$abundance_{codon} > max((33 - mean(min(Q\_score)) / 0.015, 11) \quad \text{(Eq. 1).}$$

The filter values were set to minimize the number of identified mutated codons in the sequence of the non-mutated AIDA autotransporter adjacent to the hArg1 gene. This process produced a variant table that identified in-frame codon substitutions and linked them to specific read-IDs and experimental bins.

**Amber-Scanning Fitness Score Calculation.**

To represent and compare different variants, a fitness score was calculated for each variant and each experiment. In the case of SAC incorporation, the fitness score was a representation of the weighted fluorescence mean of each variant for display labeling (i.e. expression fitness score). For the click chemistry reactions, the fitness score was calculated from a weighted mean of the fluorescence intensity derived from bioconjugation with the fluorophore or the PEG molecule.

To do so, we estimated the number of cells ($C_{vi}$) in each bin ($i$) of a specific variant ($v$) by assuming the number of reads is representative of the underlying cell population and multiplying the abundance of a variant by the total number of sorted cells (Equation 2).

$$C_{vi} = \frac{reads\ variant_i}{reads\ total_i} \times total\ cells\ in\ bin_i \qquad \text{(Eq. 2).}$$

To improve reliability and reduce bias from sequencing errors, low-occurring variant reads were filtered out to retain variants with least $C_{vi} > 15$. For each biological replicate ($rep_j$), a weighted mean across bins was calculated using the median fluorescence value ($med\_fluor_i$) as the weight (Equation 3). The median fluorescence was calculated from a sample recording during each sort.

$$fitness_{rep\_j} = \frac{\sum_{i=1}^{4} C_{vi} \times med\_fluor_i}{\sum_{i=1}^{4} C_{vi}} \qquad \text{(Eq. 3).}$$

Finally, the base-10 logarithm was taken of these values (Equation 4) and then a second weighted mean was calculated among the replicates for the final fitness value per variant (Equation 5).

$$log\_fitness_{rep\_j} = log_{10}(fitness_{rep\_j}) \qquad \text{(Eq. 4).}$$

$$final\_fitness = \frac{\sum_{j=1}^{2} log\_fitness_j \times C_{vi}}{\sum_{j=1}^{2} C_{vi}} \qquad \text{(Eq. 5).}$$

**Calculation and Data Collation of hArg1 Physical Properties.**

To understand the effect of hArg1 physical properties on fitness scores, we collected and calculated a series of physical properties of hArg1. The relative solvent-accessible surface area (relSASA) of each residue side chain for monomeric hArg1 (PDB: 6Q92) was calculated using the get_relative_sasa function in PyMol (https://pymolwiki.org/index.php/Get_sasa_relative). B-factors were extracted from the same PDB file. Statistical comparisons were performed using ANOVA or Kolmogorov-Smirnov (KS) tests, with normality analyzed. ANOVA was used for normally distributed data, otherwise, the KS test was performed.

**PEGylation reaction for epitope mapping.**

To map the epitope of an anti-hArg1 antibody, SAC-incorporated hArg1 displaying cells were bioconjugated with 15 µM metz-$PEG_{23}$-DBCO (BroadPharm, BP-25743) and 30 µM azide-AF594 (Jena Bioscience, CLK-1295-1) following the amber-scanning experiment procedure with an extended incubation of 40.5 hours. Cells were washed twice with PBS-BSA-T20 and twice with PBS-BSA before performing a immunofluorescence labeling assay with the mouse anti-hArg1 antibody (Invitrogen, MA5-24298). 200 million cells were incubated with a 1:15,000 dilution of the antibody at room temperature for 30 minutes. After two washes in PBS-BSA, they were incubated in ice for 20 minutes with a 1:500 dilution of goat anti-mouse IgG AF488. Labeled cells were kept at 4 ºC until analyzed by flow cytometry in a BD FACSMelody.

**Flow cytometry and ONT sequencing for epitope mapping.**

Immunolabeled and PEGylated cells were sorted in three bins in biological replicates and at least 1,500,000 cells were collected per replicate. Similarly to the amber-scanning library, DNA was purified and cleaned. The hArg1 gene was extracted by restriction digestion and then barcoded and indexed for ONT sequencing following the same procedure as described above. Basecalling was performed on Dorado (version 0.5.2+7969fab) using the dna_r10.4.1_e8.2_400bps_sup@v4.3.0 basecalling model, and variant calling was performed using the same strategy as before.The process resulted in >2.2 million number of filtered reads.

To calculate the variant abundance in each bin, we firstly estimated the number of cells per bin and variant ($C_{vi}$), as before (Equation 1). Then, we calculated the abundance of every variant per bin (Equation 6).

$$abundance\_bin_i = \frac{C_{vi}}{C_{v1}+C_{v2}+C_{v3}} \quad \text{(Eq. 6).}$$

Highly abundant variants in Bin 1 ($value_{bin\ 1} > mean_{bin\ 1} + 1.5\ SD_{bin\ 1}$) were selected excluding very highly-abundant variants in Bin 2 ($value_{bin\ 2} < mean_{bin\ 2} + 2\ SD_{bin\ 2}$). The identified residues were represented in red as sticks on a trimeric representation of hArg1 (from PDB: 6Q92).

**Single variant $K_D$ measurements**

For the display of the hArg1 single variants, the constructs were transformed into *E. coli* BL21(DE3) cells and SAC was incorporated as described above in autoinduction media supplemented with 4 mM SAC, appropriate antibiotics and grown for 22-24h. The $OD_{600}$ was adjusted with PBS to 2.5 (2.0 x $10^9$ cells/mL) with a final volume of 1 mL. These cells were washed twice in PBS and again adjusted to 1 mL volume. Then the 1 mL stock was split and transferred to a 13 mL tube. To one half of the samples metz-$PEG_{23}$-DBCO (BroadPharm, BP-25743) was added to a final concentration of 30 µM. All samples were incubated at 7 °C for 48h, with shaking at 240 rpm. These 500 µL stocks were then split in 10 x 50 µL (1.0 x $10^8$ cells each) and placed into 10 wells of a 96-well plate. The cells were washed twice with PBS-T20 and once with PBS (100 µL buffer volume, centrifuged for 5 min at 2204 x g, supernatant discarded). Afterwards the cells were resuspended in 50 µL of the dilution series buffer (PBS) containing the hArg1 antibody (Invitrogen, MA5-24298) and incubated for 30 min. The dilution series started from 1:50 with a dilution factor of 3 and 9 dilution steps, covering the concentration range from 66.67 nM to 0.0034 nM. Afterwards, the cells were washed with PBS and resuspended in PBS containing a secondary AF488 conjugated antibody. The cells were incubated for 30 min, washed with PBS, then 1:10 diluted in the PBS and measured by flow cytometry with a Attune NxT flow cytometer (ThermoFisher Scientific) to generate a titration curve.

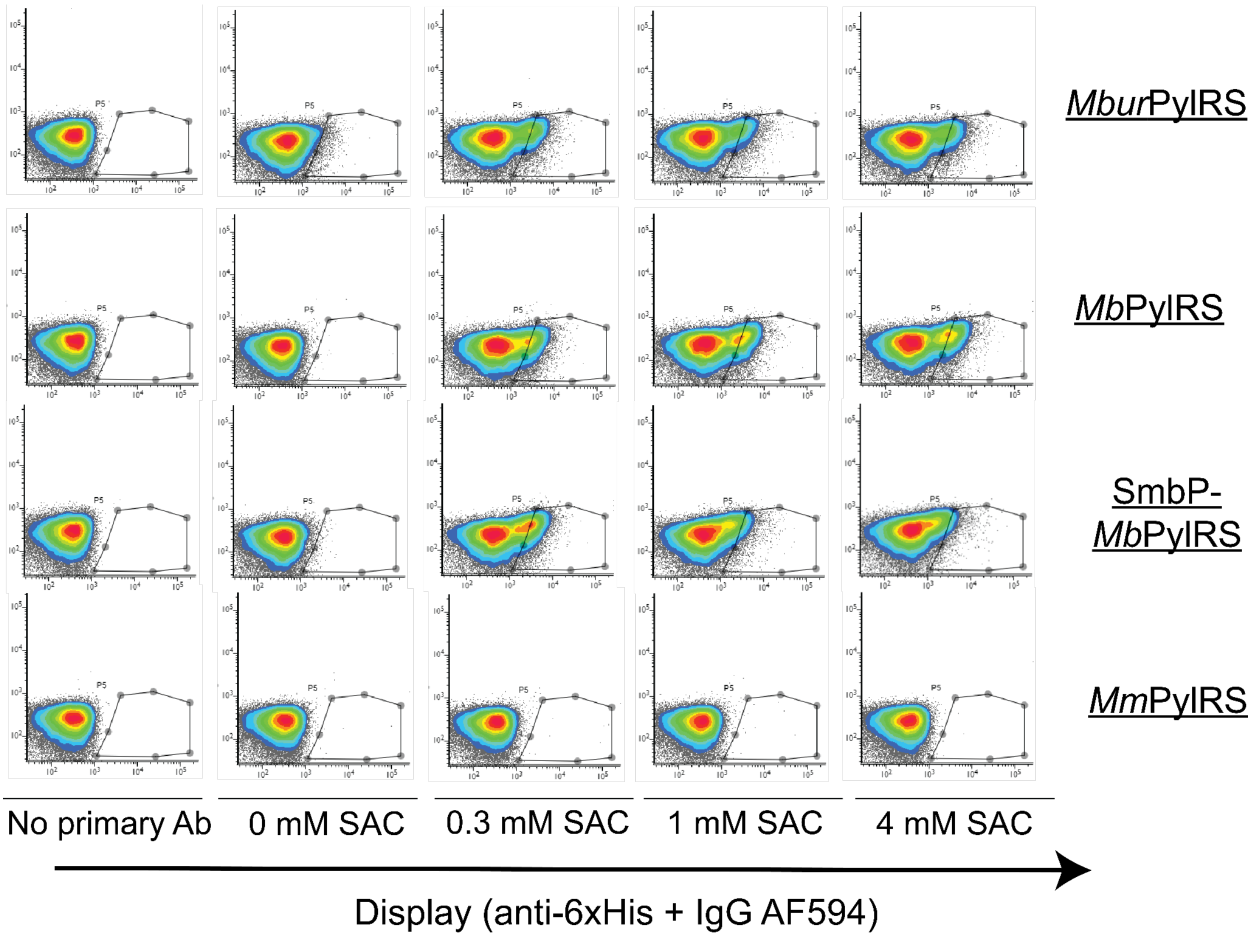


**Figure S1: PylRS comparison for SAC incorporation efficiency on displayed hArg1.**
Immunofluorescence assay characterizing hArg1 display level in the presence of amber-stop codons with different bioorthogonal translational systems. Each dot plot shows a representative sample from overall biological triplicates.

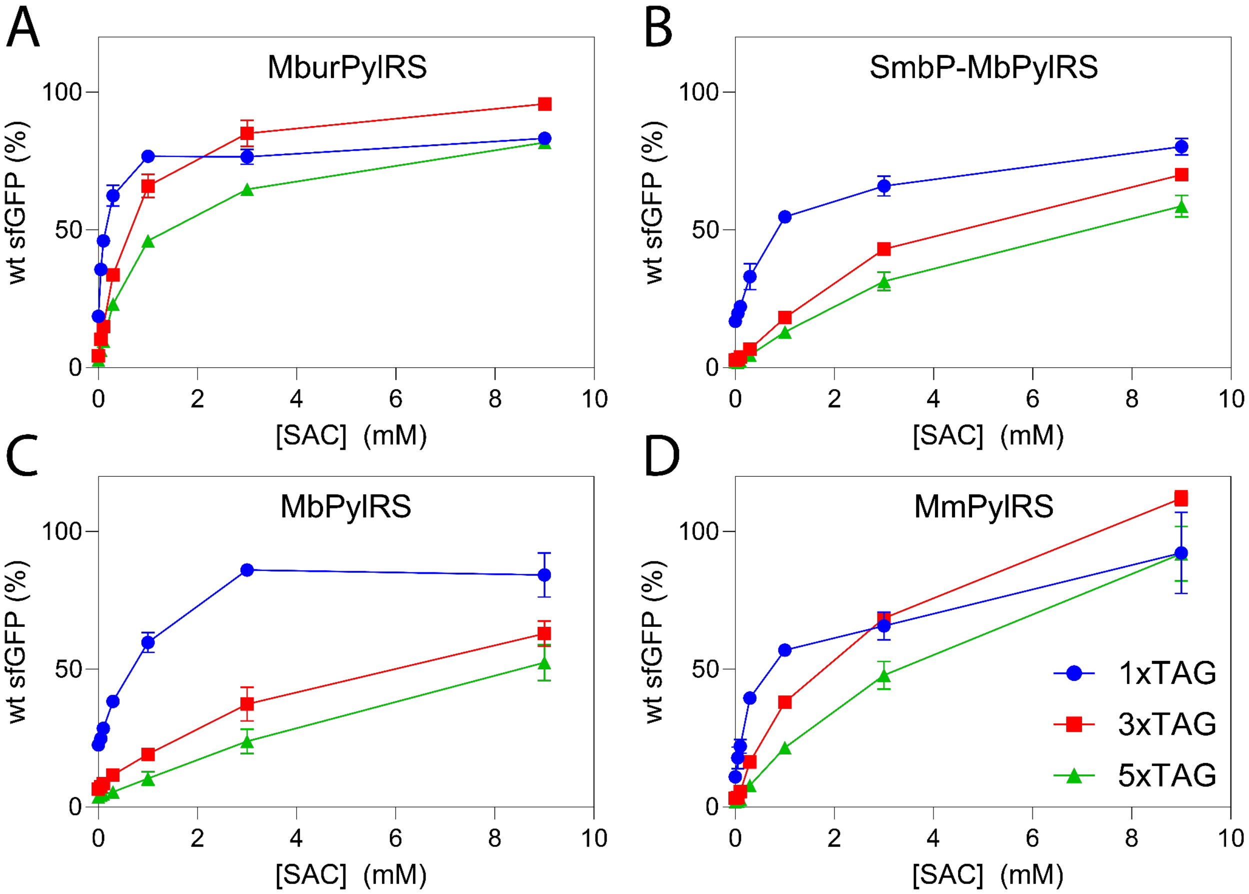


**Figure S2**: **Reporter assay with sfGFP.**

Reporter assay for ncAA incorporation efficiency at a sfGFP with 1x (blue), 3x (red) or 5x (green) amber stop codons at different SAC concentrations. (A) *Mbur*PylRS, (B) SmbP-*Mb*PylRS, (C) *Mb*PylRS and (D) *Mm*PylRS were compared in a *E. coli* strain optimized for GCE (B-95.ΔAΔfabR).[6] Data points represent the mean percent of activity compared to wildtype sfGFP and the error bars represent the SD of three biological replicates.

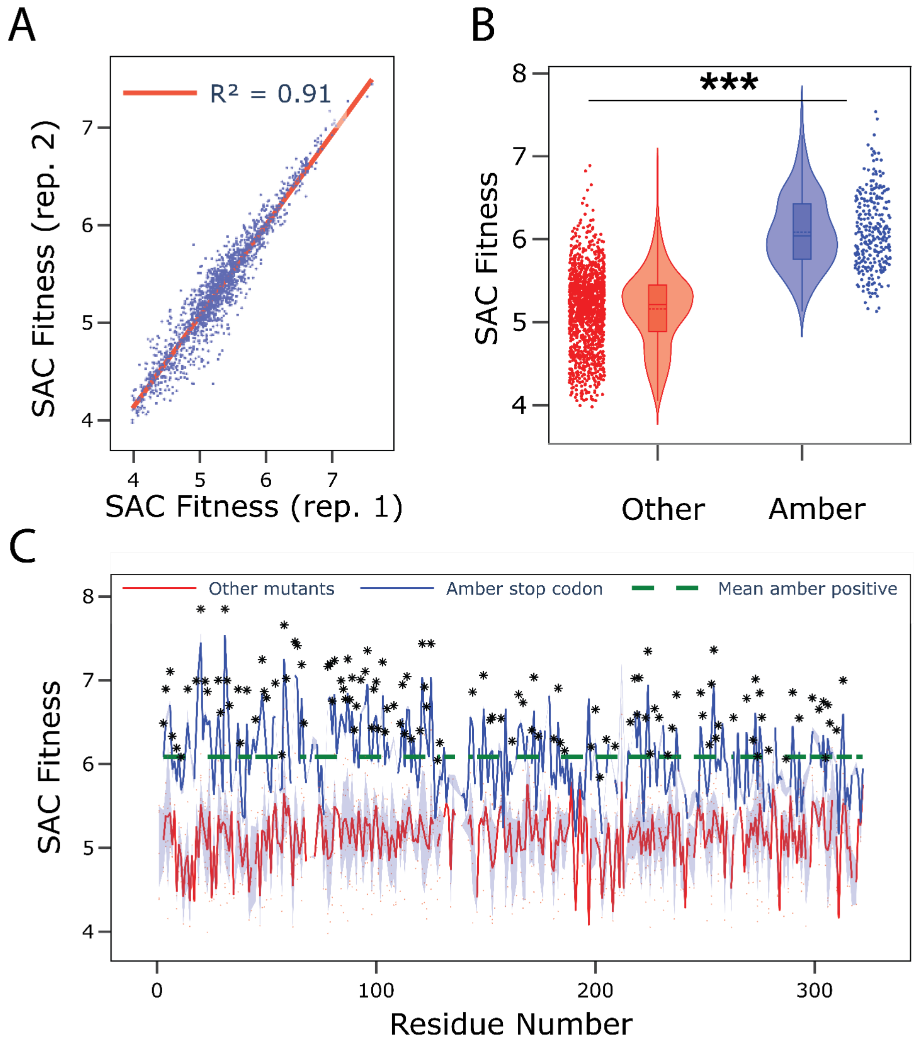


**Figure S3: SAC amber-scanning validation.**

(A) SAC fitness score reproducibility among biological replicates. The dataset was filtered for variants carrying single amber stop codons and represented by at least 15 cells in the dataset ($C_{vi} \geq 15$). The Pearson's correlation coefficient ($R^2$) is given. (B) Distribution of fitness scores for amber stop codons in comparison to any other mutant for the incorporation of SAC. (*** represents a p-value < 0.001 for both a Student t-test and a Kolmogorov-Smirnov (KS) test). (C) SAC fitness score per residue number for amber stop codons (blue) and other variants (red) where the shaded areas represent the SD. The green dashed line the average fitness across amber positive population. * represent a p-value < 0.05 for a KS-test.

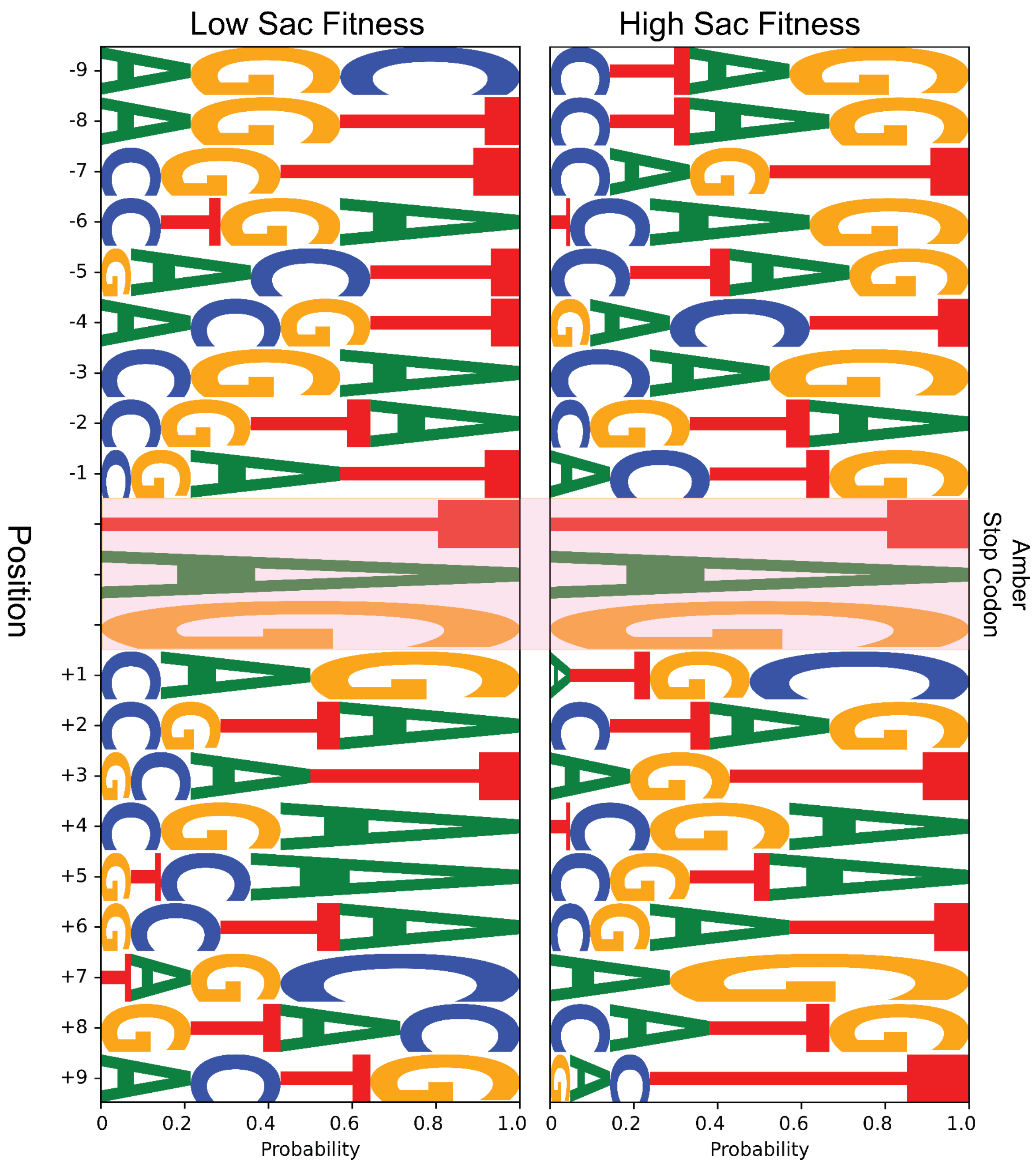


**Figure S4: Incorporation efficiency of SAC by nucleotide context.**

Nucleotide preference around amber stop codon for high (value > mean + 1.5*SD) (left) and low (value < mean - 1.5*SD) (right) SAC fitness scores. n= -1 is the nucleotide directly upstream of the amber-stop codon and n = +1 is the one directly downstream.

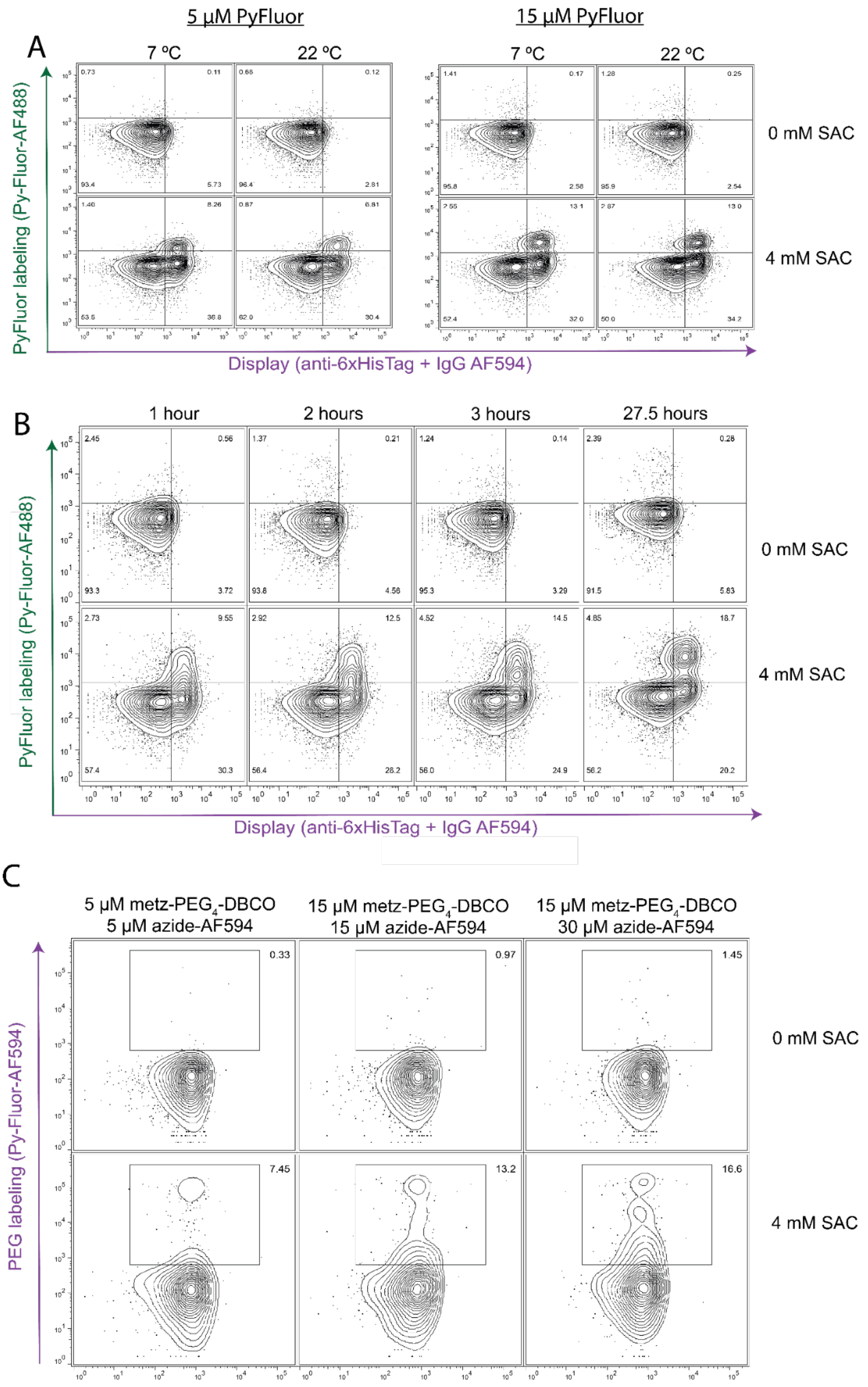


**Figure S5: Fluorophore and PEG reaction conditions optimization.**
(A) Temperature and concentration optimization of fluorophore binding. (B) Time course for fluorophore binding. (C) Concentration of reactants for PEGylation.

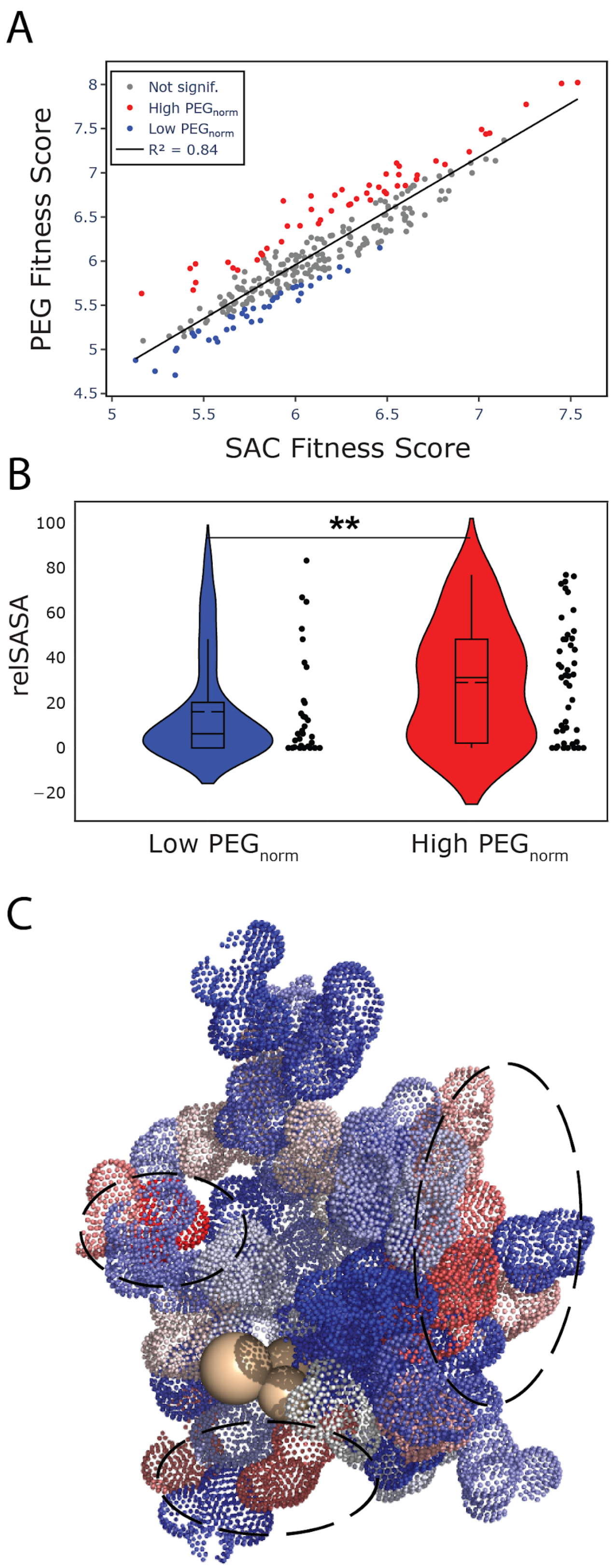


**Figure S6: Dynamics of hArg1 studied by PEG amber-scanning.**

Filtered variants according to high (red) or low (blue) $PEG_{norm}$. Variants within one SD of the mean were considered as not significant (n.s.). (B) Density distribution of relSASA values for low and high solvent accessible variants. (** for p-value < 0.01, KS-test). (C) Surface representation of fully buried residues (relSASA = 0) with color representing PEGnorm values. The darkest blues represent low PEGnorm values and the darkest red represents the highest PEGnorm values.

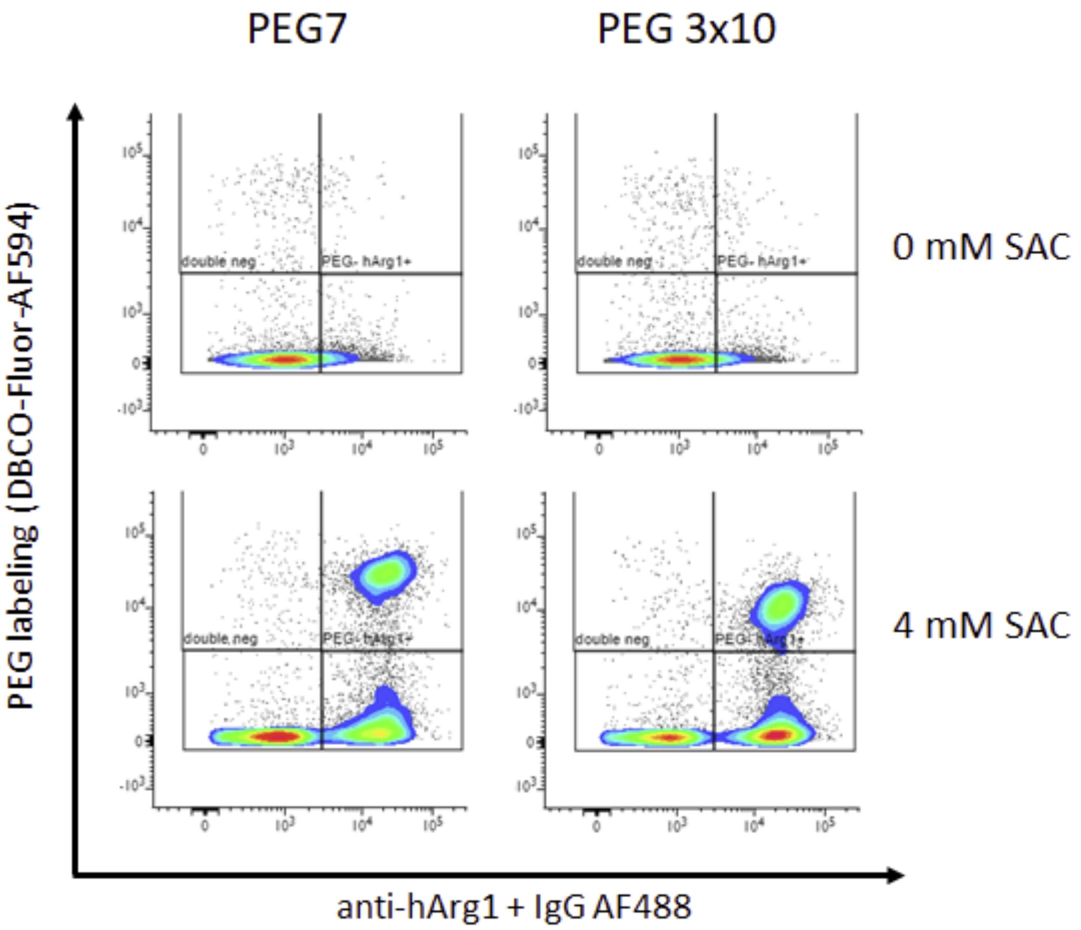


**Figure S7: Fluorophore and PEG reaction with PEG7 and branched PEG 3x10.**
Bioconjugation of displayed hArg1 with Methyltetrazine-amido-PEG7-azide (BroadPharm catalog # BP-23552) and a branched PEG consisting of tri-branched PEG10, (Methyltetrazine-PEG10)-Tri-(Azide-PEG10-ethoxymethyl)-methane (BroadPharm catalog # BP-23552). The fluorophore was DBCO-AZDye594 (Jena Bioscience catalog # CLK-1298-AZ-1).

**Table S1. Statistical analysis of binding affinity of selected variants tested as monogenic cultures.** Statistical analysis was performed using one-way-ANOVA Brown–Forsythe test with a consecutive Dunnett's T3 multiple comparisons test. For the N319 mutant, no $K_D$ (i.e. no binding) was detected. For the statistical analysis, we used an arbitrarily high value (mean of 140 nM) for the $K_D$ to simulate the significance test.

| | **PEGylated** | | **SAC** | |
|---|---|---|---|---|
| | Comparison | Adjusted P-value | Comparison | Adjusted P-value |
| **epitope** | wt:PEG vs D181:PEG | 0.0159 | wt:SAC vs D181:SAC | 0.0189 |
| | wt:PEG vs E309:PEG | 0.0194 | wt:SAC vs E309:SAC | 0.1748 |
| | wt:PEG vs G305:PEG | 0.0528 | wt:SAC vs G305:SAC | 0.0879 |
| | wt:PEG vs P157:PEG | 0.0528 | wt:SAC vs P157:SAC | 0.0716 |
| | wt:PEG vs P184:PEG | 0.0062 | wt:SAC vs P184:SAC | 0.0013 |
| | wt:PEG vs Q43:PEG | 0.001 | wt:SAC vs Q43:SAC | 0.216 |
| | wt:PEG vs T136:PEG | 0.0227 | wt:SAC vs T136:SAC | 0.2234 |
| | wt:PEG vs T295:PEG | 0.0382 | wt:SAC vs T295:SAC | 0.1935 |
| | wt:PEG vs Y317:PEG | 0.1296 | wt:SAC vs Y317:SAC | 0.0024 |
| | wt:PEG vs N319:PEG | <0.0078 | wt:SAC vs Y319:SAC | 0.0077 |
| **non-epitope** | wt:PEG vs A56:PEG | 0.9992 | wt:SAC vs A56:SAC | 0.2435 |
| | wt:PEG vs I58:PEG | 0.1413 | wt:SAC vs I58:SAC | 0.9115 |
| | wt:PEG vs K68:PEG | 0.9841 | wt:SAC vs K68:SAC | 0.1475 |
| | wt:PEG vs S109:PEG | 0.7662 | wt:SAC vs S109:SAC | 0.6532 |
| | wt:PEG vs S62:PEG | 0.1464 | wt:SAC vs S62:SAC | 0.9817 |

**References.**
